# Quadratization: From conductance-based models to caricature models with parabolic nonlinearities

**DOI:** 10.1101/137422

**Authors:** Axel G. R. Turnquist, Horacio G. Rotstein

## Abstract

*Quadratization* of biophysical (conductance-based) models having a parabolic-like voltage nullcline in the subthreshold voltage regime refers to the process by which these models are substituted by “caricature” models having a strictly parabolic voltage nullcline and a linear nullcline for the recovery variable. We refer to the latter as quadratic or parabolic models. The parabolic-like and strictly parabolic voltage nullclines coincide at their extrema (minima or maxima) and are well approximated by each other in vicinities of these extrema whose size depend on the model parameters. Quadratic models are simplified by a change of variables that translates these extrema into the origin of the phase-plane diagram. A further simplification (parameter reduction) can be achieved by nondimensionalizing the quadratic models. This procedure can be extended to three-dimensional models having a parabolic-cylinder-like shaped voltage nullsurface and to models having time-dependent inputs and synaptic currents.

## Detailed Description Introduction

### Introduction

Reduced models of neurons often involve a detailed description of the subthreshold dynamics supplemented with artificial spikes when the variable representing the voltage (*V*) crosses a preset threshold and additional reset values for all or some of the participating variables after a spike has occurred.

The different types of mathematical description of the subthreshold dynamics can be broadly classified into two groups: biophysical and phenomenological. Biophysical models include the description of the interplay of the ionic currents that operate in the subthreshold voltage regime. These models can be thought of as conductance-based models of Hodgkin-Huxley (HH) type (Hodgkin and Huxley 1952) where the spiking currents (transient sodium and delayed-rectifier potassium) have been removed (Lapicque 1907; Ermentrout and Terman 2010; Gerstner et al. 2014; Rotstein et al. 2006; Smith et al. 2000; Izhikevich 2006). Phenomenological models consist of heuristic dynamic equations that include the types of nonlinearities and time constants that are assumed to be present in the system and determine the geometry of the phase-space diagrams. The parameters have no explicit connection to the neuronal biophysics, but they can be fitted to the observed experimental patterns and therefore these models can be used to understand neuronal dynamic mechanisms in terms of relatively simple sets of equations. Some models lie in between these two groups.

In the so-called leaky integrate-and-fire (IF) model the subthreshold dynamics are linear and one-dimensional. The quadratic and exponential IF models have also one-dimensional subthreshold dynamics and include the said type of nonlinearities for the evolution of the equation for *V* (Latham et al. 2000; Hansel and Mato 2001; Fourcaud-Trocme et al. 2003). Models with two-dimensional dynamics can be obtained by adding a recovery variable (*w*) with linear dynamics to the models mentioned above, giving rise to the linear resonate-and-fire models (Izhikevich 2001) and the non-linear adaptive quadratic (Izhikevich 2003) and exponential (Brette and Gerstner 2005) IF models. More complex models with three-dimensional subthreshold dynamics involve two recovery variables corresponding to ionic currents with slow dynamics (e.g., hyperpolarization-activated mixed sodium-potassium, T-type calcium, adaptive) (Rotstein et al. 2006; Smith et al. 2000).

Phenomenological models of quadratic IF type with strictly parabolic *V*-nullclines in the sub-threshold regime (Izhikevich 2003) have been frequently used as ad-hoc models to investigate the dynamics of neurons in a variety of systems. The lack of connection with the neuronal biophysics limits the scope of these studies as well as network studies that combine the two modeling approaches (phenomenological and biophysical) for different neuron types in the same circuits (Tikidji-Hamburyan et al. 2015; David et al. 2015). For example, phenomenological models cannot capture the effects of changes in the biophysical conditions (e.g., ionic currents levels, synaptic connectivity) on the resulting patterns. For yet another example, these models also fail to capture the existence of patterns with equal attribute values (e.g., firing frequency) generated by neurons with different biophysical properties (Prinz et al. 2004; Rotstein et al. 2016) or the effects of these types of heterogeneous neurons on the network patterns in which they are embedded. These issues are not unique for models of adaptive quadratic IF models, but they are shared by all phenomenological models. Linearization of conductance-based models (Richardson et al. 2003; Koch 1999) provides explicit formulas for the parameters in the linearized models in terms of the biophysical parameters (Izhikevich 2001; Rotstein 2013, 2014), but linear models fail to capture important dynamic effects.

In this paper we present a systematic reduction of biophysical (conductance-based) models with parabolic-like *V*-nullclines (nullsurfaces) to phenomenological models with strictly parabolic *V*- nullclines (nullsurfaces) and linear nullclines (nullsurfaces) for the recovery variable(s). We refer to these reduced models as parabolic or quadratic and to the reduction process as quadratization. The approach presented here extends the concept of linearization to include second order derivatives in voltage and captures the quadratic nonlinearities in close vicinities of the extrema of the parabolic-like nullclines for the underlying two-dimensional system.

We focus on two-and three-dimensional models having one or two voltage-dependent gating variables, respectively (each of which corresponding to an ionic current) and an additional gating variable with fast dynamic (slave to voltage). The system consisting of the voltage equation together with an equation for one of the gating variables (*x*_1_) is thought to be the “main” system. The second gating variable (*x*_2_), typically slower, is considered to have a modulatory effect in the sense that the projections of the *V*-nullsurface onto planes parallel to the *V*-*x*_1_ plane remain parabolic-like in the subthreshold regime. We also include a time-dependent input current and a synaptic current in order to make the reduction amenable for networks models.

After an appropriate translation, the minimum of the strictly parabolic *V*-nullcline is located at the origin of the phase-plane diagram. The strictly parabolic *V*-nullcline is concave up. If the parabolic-like *V*-nullcline is concave-down, then the quadratization process includes an inversion of the *V*-nullcline. An additional simplification (nondimensionalization) reduces the number of parameters further, but it changes the definitions of the time and the dependent variables. Importantly, the units of the variables in the quadratic model are different from their original units. This approach produces formulas that relate the parameters of the quadratic model with the biophysical parameter. This is necessary for the biophysical interpretation of the results using caricature models.

The quadratization procedure described in this paper assumes that the *V*-nullcline (nullsurface) is parabolic-like (parabolic-cylinder-like shaped) in the subthreshold voltage regime. This is often overlooked when phenomenological models of quadratic IF type with strictly parabolic *V*-nullclines are used. In fact, the neuronal subthreshold dynamics is often governed by other types of nonlinearities such as cubic-like ones (Rotstein 2017a,b; Remme et al. 2012). Whether the subthreshold dynamics have parabolic-or cubic-like nonlinearities may not be apparent from the identity of the participating ionic currents in the biophysical models (Izhikevich 2006; Rotstein 2017a). As shown in previous work, models with the same combination of ionic currents may give rise to different types of nonlinearities (parabolic-and cubic-like) in the subthreshold regime (Rotstein 2017a).

We discussed additional limitations of the quadratization procedure and the use of quadratic models later in the paper.

## Models

### Conductance-based models

We use conductance-based models of Hodgkin-Huxley type (Hodgkin and Huxley 1952) to describe the neuronal subthreshold dynamics. Our formulation allows for various combinations of ionic currents that have been shown to underlie the generation of subthreshold resonance, including persistent sodium (*I*_*Nap*_), h-(*I*_*h*_) and slow potassium currents (*I*_*Ks*_) (Acker et al. 2003; Schreiber et al. 2004; Rotstein et al. 2006; Izhikevich 2006) (see below). However, we do not include the spiking currents (transient sodium and delayed-rectifier potassium). The onset of spikes is described by the models we use. However, in order to describe the spiking dynamics either these currents have to be included or spikes have to added manually after their onset.

For generality, in our description we use non-specific ionic currents. We discuss specific model examples later in this section. The current-balance equation is given by

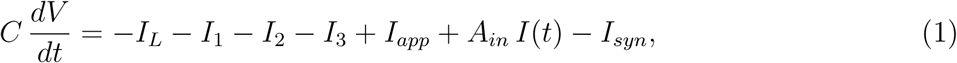

where *V* is the membrane potential (mV), *t* is time (msec), *C* is the membrane capacitance (*μ*F/cm^2^), *I*_*app*_ is the applied bias (DC) current (*μ*A/cm^2^), *I*(*t*) is a time-dependent input current with amplitude *A*_*in*_ (*μ*A/cm^2^), *I*_*L*_ = *G*_*L*_ (*V*- *E*_*L*_) is the leak current, *I*_*j*_ (*j* = 1, 2, 3) are ionic currents and *I*_*syn*_ is a synaptic current. The ionic currents we consider here are of the form

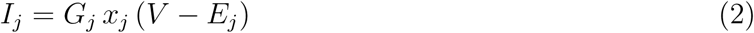

where *x*_*j*_ are the gating variables, *G*_*j*_ the maximal conductances (mS/cm^2^) and *E*_*j*_ the reversal potentials (mV).

All gating variables *x* obey a first order differential equation of the form

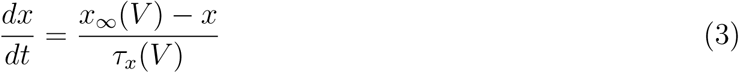

where *x*_*∞*_(*V*) and *τ*_*x*_(*V*) are the voltage-dependent activation/inactivation curves and time-scales respectively.

The synaptic current we consider has the form

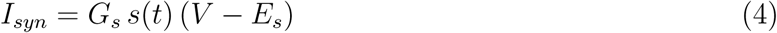

where *s*(*t*) is the synaptic function, *G*_*syn*_ the maximal synaptic conductance and *E*_*syn*_ the synaptic reversal potential.

The ionic currents we consider here (2) are restricted to have a single gating variable *x* and to be linear in *x*. In addition, in this paper we focus on two-dimensional models having two dynamic gating variables (*x*_1_ and *x*_2_) and an additional gating variable evolving on a fast time scale for which the adiabatic approximation *x*_3_ = *x*_3, ∞_(*V*) is justified. Additional fast currents can be included without significantly changing the formalism.

### Models of quadratic type

The models of quadratic type we use in this article have the form

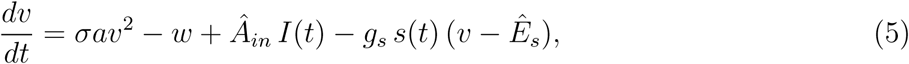

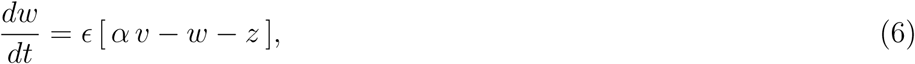

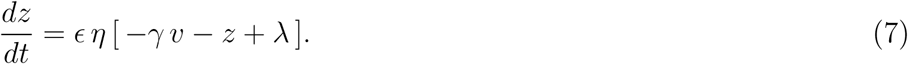

For two-dimensional biophysical models the geometric model reduces to eqs. (5)-(6) with *z* substituted by a constant (*z* = *λ*).

Collectively, we refer to the parameters *a*, *α*, *γ*, *λ* and *σ* as the geometric parameters since they control the geometry of the phase-space diagram: *α* and *γ* control the slope of the *w*_1_- and *w*_2_-nullsurfaces, respectively, *λ* controls the relative displacement between the *v*- and the *w*_2_-nullsurfaces (or between the *v*- and *w*_1_-nullclines in the two-dimensional case). The parameter *a* is assumed to be positive. The product *σ a* controls the curvature of the parabolic *V*-nullcline. The concavity sign is captured by *σ* = ±1. By an appropriate change of variables when *σ* = -1 (concave down parabolic *v*-nullcline) the model can be transformed into one having a concave up parabolic *v*-nullcline.

The dynamic parameters *∊* and *η* represent the time scale separation between the variables *v* and *w*_1_, and between the variables *w*_1_ and *w*_2_, respectively. Without loss of generality, we assume that *η <* 1 as we discuss below.

The geometric and dynamic parameters and the parameters *Â*_*in*_, *g*_*s*_ and *E*_*s*_ can be expressed in terms of the biophysical parameters in eqs. (1)-(4). We provide these expressions below and briefly explain the quadratization procedure that brings the biophysical model (1)-(4) into the quadratic model (5)-(7).

The units of the variables and parameters in eqs. (5)-(7) are [*v*] = mV, [*w*] = mV/msec, [*z*] = mV/msec, [*∊*] = 1/msec, [*η*] = 1, [*a*] = 1/(msec mV), [*α*] = 1/msec, [*γ*] = 1/msec, [*λ*] = mV/msec, 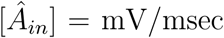, 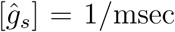 and [*E*_*s*_] = mV. Some of these variables and parameters involve the capacitance *C*. When it is assumed to be *C* = 1 and not included in the definitions of the variables and parameters, [*w*] = mS mV/cm^2^, *z* =, mS mV/cm^2^, [*a*] = mS/(cm^2^ mV), [*α*] = mS/cm^2^, [*γ*] = mS/cm^2^, [*λ*] = mS mV/cm^2^, 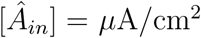 and 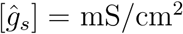,

### Dimensionless models of quadratic type

System (5)-(7) can be simplified to the following model with a reduced number of parameters

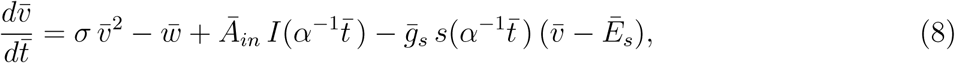

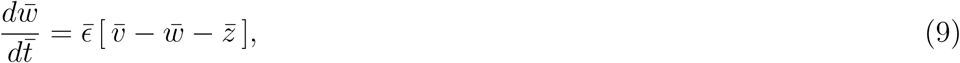

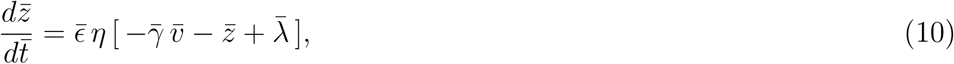

by a nondimensionalization process, where

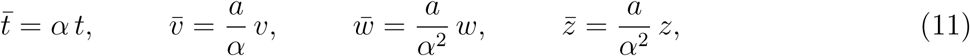

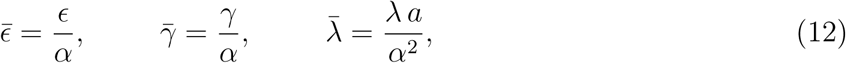

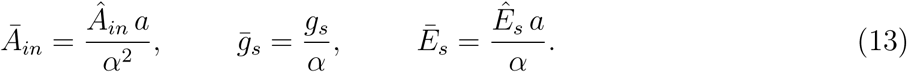

## Quadratization procedure

Without loss of generality, we assume that *τ*_2_ < *τ*_1_ (*x*_2_ is slower than *x*_1_). We also assume that the projection of the *V*-nullsurface onto the planes parallel to the *V*-*x*_1_ plane (constant values of *x*_2_) are parabolic-like functions. In other words, we assume that the presence of the variable *x*_2_ does not qualitatively change the shape of the parabolic-like *V*-nullcline of the two-dimensional system (for *x*_2_ = 0), but it can only produce parabolic-like deformations, raise it and shift it down as in Rotstein et al. (2006). The case *τ*_2_ = *τ*_1_ is excluded because the system can be reduced to a quasi-two-dimensional (Rotstein 2017a).

As a starting point we identify the minimum (*V*_*min*_, *x*_1,*min*_, 0) or maximum (*V*_*max*_, *x*_1,*max*_, 0) of the parabolic-like *V*-nullcline (for *x*_2_ = 0). We refer to them indistinctly as (*V*_*e*_, *x*_1,*e*_, *x*_2,*e*_). We define

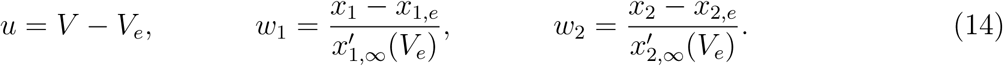

We then expand the right-hand sides of (1) and (3) around (*V*_*e*_, *x*_1,*e*_, *x*_2,*e*_) up to the first order in *x*_1_ and *x*_2_ and up to the second order in *V*. This yields

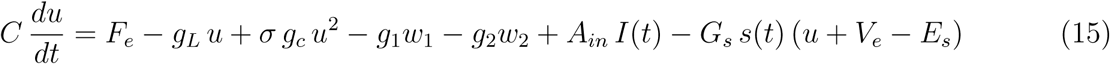

and

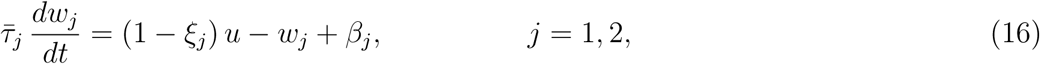

where

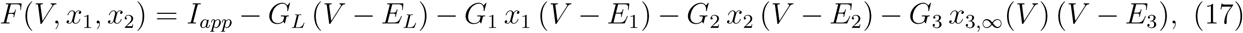

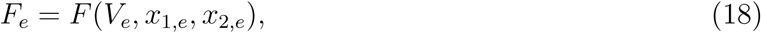

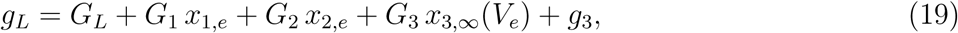

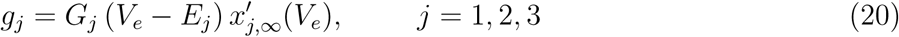

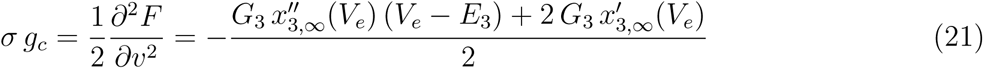

and

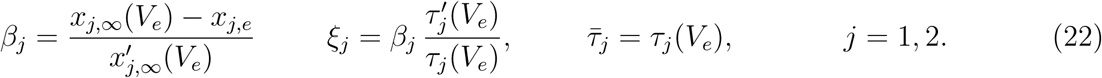

In (21) *g*_*c*_ > 0 and the concavity of the original *V*-nullcline is captured by the sign of *σ* = ±1.

Next, we define

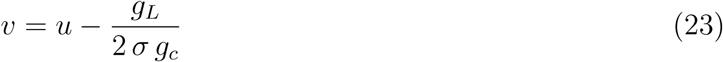

Substituting into (15)-(16) yields

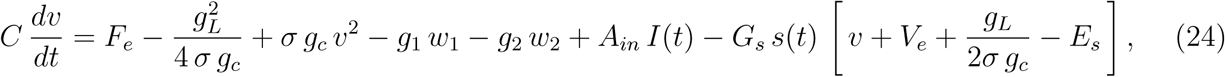

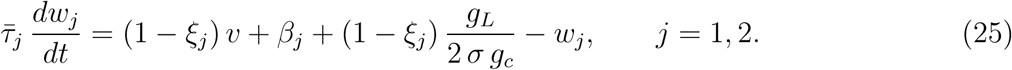

Subsequently, we define

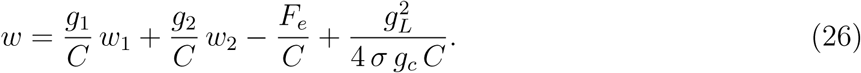

Substituting into (24)-(25) we obtain

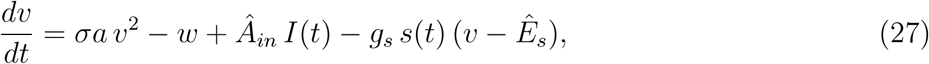

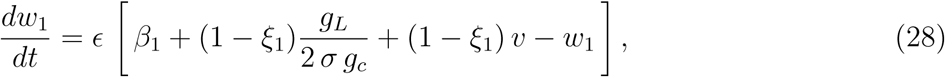

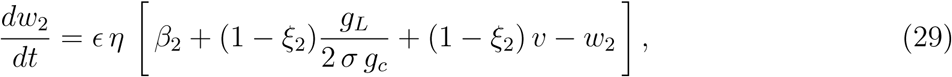

where

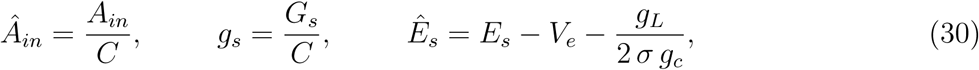

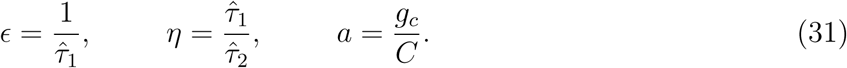

By differentiating both sides of eq. (26) with respect to *t*, substituting (28) and (29) and rearranging terms we obtain eq. (6) where

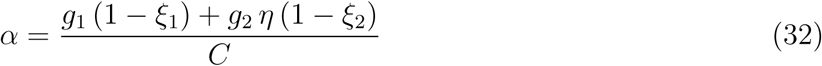

and

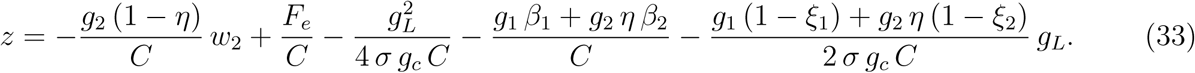

By differentiating both sides of eq. (33) with respect to *t*, substituting (29) and rearranging terms we obtain eq. (7) where

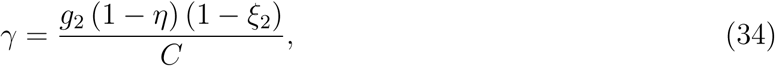

and

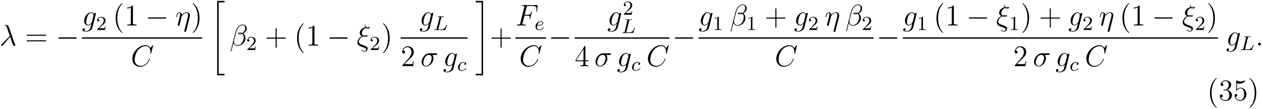

For *g*_2_ = 0, substituting into (31), (32) and (31) yields *γ* = 0 and *z* = *λ*.

## Two examples

We present two two-dimensional and autonomous (*G*_2_ = *G*_*s*_ = *A*_*in*_ = 0 and *g*_2_ = *g*_*s*_ = *Â*_*in*_ = 0) examples: the *I*_*Nap*_ + *I*_*h*_ and *I*_*Nap*_ + *I*_*Ks*_ models (Rotstein 2017a, 2015) adapted from (Acker et al. 2003; Rotstein et al. 2006).

The *I_Nap_* + *I_h_* involves the interaction between a persistent sodium (*I*_*Nap*_ = *I*_3_) and a hyperpolarization-activated (or h-) (*I*_*h*_ = *I*_1_) currents, where *I*_*Nap*_ = *G*_*p*_ *p*_*∞*_(*V*) (*V* – *E*_*Na*_) and *I*_*h*_ = *G*_*h*_ *r* (*V* – *E*_*h*_). The *I*_*Nap*_ + *I*_*Ks*_ model involves the interaction between a persistent sodium (*I*_*Nap*_ = *I*_3_) and slow potassium (*I*_*Ks*_ = *I*_1_) currents, where *I*_*Nap*_ = *G*_*p*_ *p*_*∞*_(*V*) (*V* – *E*_*Na*_) and *I*_*Ks*_ = *G*_*q*_ *q* (*V* – *E*_*k*_). We use the same *I*_*Nap*_ in both models. The activation and inactivation curves we used are given by the following functions (see Fig. 1)

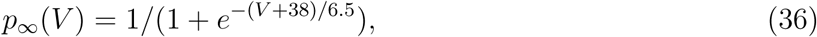

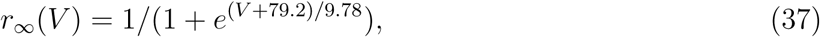

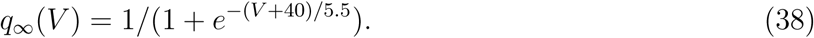

**Figure 1:**
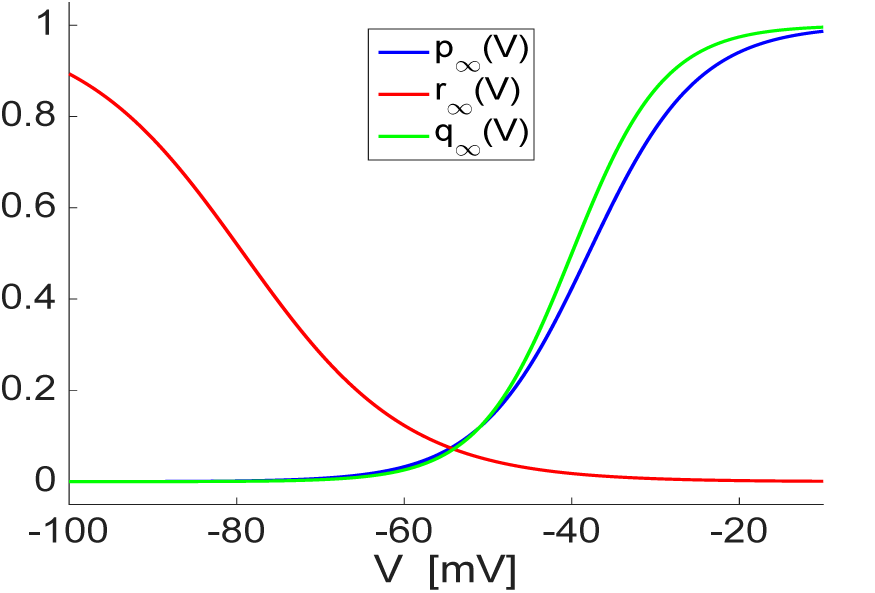
Activation/inactivation curves for the *I*_*Nap*_+*I*_*h*_ and *I*_*Nap*_ + *I*_*Ks*_ models. The function *p*_*∞*_(*V*), *r*_*∞*_ (*V*) and *q*_*∞*_(*V*) are given by (36), (37) and (38).

We used the following time constants *τ*_*r*_(*V*) = *τ*_*q*_(*V*) = 80. The nullclines for both models are shown in Fig. 2.

**Figure 2:**
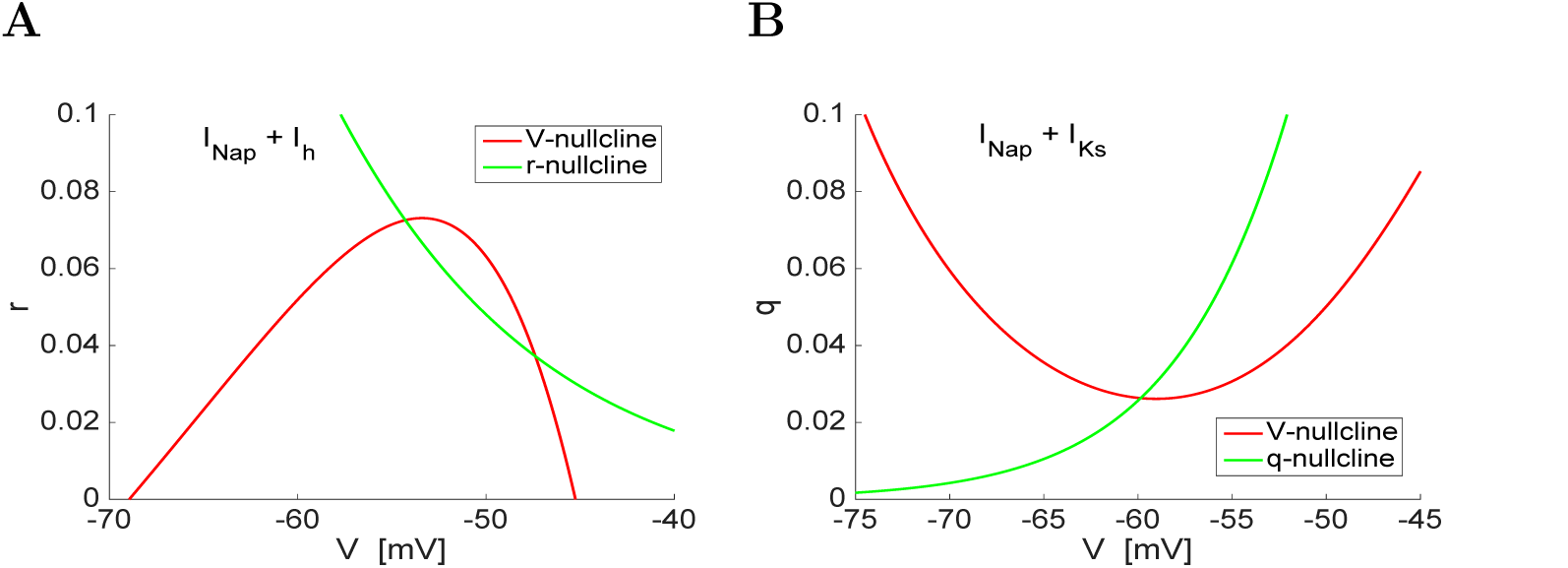
Phase-plane diagram for the 2D biophysical *I*_*Nap*_ + *I*_*h*_ and *I*_*Nap*_ + *I*_*Ks*_ models. (**A**) *I*_*Nap*_ + *I*_*h*_ model. (**B**) *I*_*Nap*_ + *I*_*Ks*_ model. The activation and inactivation curves are as in Fig. 1. We used the following parameter values (A): *C* = 1, *G*_*L*_ = 0.5, *G*_*h*_ = 1.5, *E*_*L*_ = −65, *E*_*Na*_ = 55, *E*_*h*_ = −20, *I*_*app*_ = −2.5. (B): *C* = 1, *G*_*L*_ = 0.1, *G*_*q*_ = 2, *E*_*L*_ = −54, *E*_*Na*_ = 55, *E*_*K*_ = −90, *I*_*app*_ = −0.6.

Fig. 3 shows the quadratic approximations for both models in neighborhood of the knee of the corresponding *V*-nullclines (subthreshold regime) and the linearization of the corresponding *x*_1_-nullclines (*r*- and *q*-nullclines, respectively).

**Figure 3:**
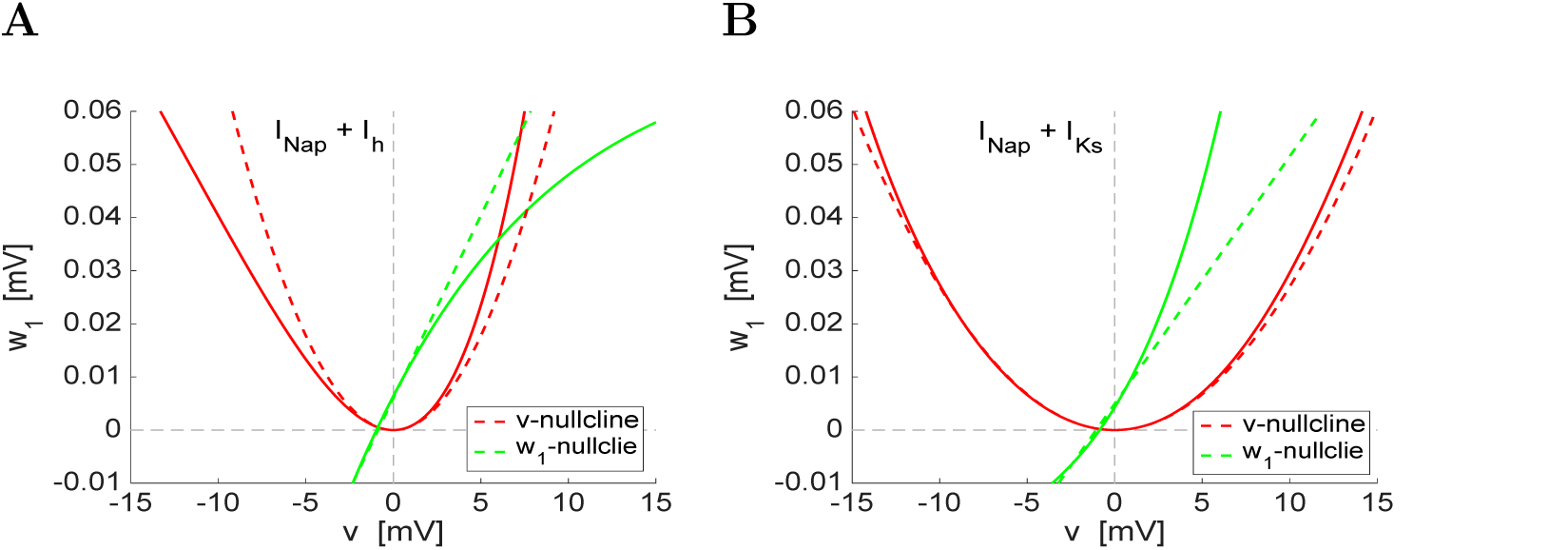
Phase-plane diagram for the 2D biophysical and parabolic (quadratized) *I*_*Nap*_ + *I*_*h*_ and *I*_*Nap*_ + *I*_*Ks*_ models. **(A)** *I*_*Nap*_ + *I*_*h*_ model. **(B)** *I*_*Nap*_ + *I*_*Ks*_ model. The solid and dashed curves are the nullclines for the biophysical (as in Fig. 2) and parabolic (quadratized) models, respectively. We used the same parameter values as in Fig. 2.

## Limitations and comments

The quadratic models (5)-(7) and (8)-(10) provide a good approximation to the original biophysical models (1)-(2) for a range of voltage values close enough to the knee of the *V*-nullcline. The parabolic-like and strictly parabolic *V*-nullclines (for *x*_2_ = 0) coincide at their extrema (minima or maxima) and are well approximated by each other in vicinities of these extrema whose size depend on the model parameters. This requires the fixed-point to be close to t the extremum (minimum or maximum) of the parabolic-like *V*-nullcline. Away from these ranges of values of *V*, but still within the subthreshold voltage regime, there may be significant differences between the parabolic-like and strictly parabolic nullclines. This requires checking the validity of the use of quadratic models in these regions where the approximations are no longer mathematically valid. One possible solution is to include higher order terms in the expansion of the right-hand side of eq. (1). However, this involves additional terms corresponding to the mixed derivatives.

In the quadratic models the recovery variable nullclines (nullsurfaces) are linear. This may create dynamic differences between the biophysical and phenomenological models. For example, the linear and parabolic nullclines may intersect twice in the subthreshold regime, while the original nullclines do not. This is more likely to happened in models with concave-up parabolic-like nullclines (e.g., Fig. 3-B) than in models with concave-down parabolic-like nullclines (e.g., Fig. 3-A). As before, one possible solution is to include second order terms in the expansion of the right-hand side of eq. (3).

When the fixed-point of the biophysical model is not close enough to the extremum of the *V*-nullcline, the biophysical model is quasi-linear in the subthreshold regime. However, the parabolic-like nonlinearities may still play a role in the global subthreshold dynamics and should not be disregarded. One possible approach in these cases is to linearize eq. (3) around the fixed-point and not around the extremum of the parabolic-like *V*-nullcline.

The use of the dimensionless model (8)-(10) involves the rescaling of *t* and *v* in addition to the other variables. This makes the comparison among results for different parameter values more difficult and less straightforward, since both *v* and *t* may be used in the computation of these attributes. This is particularly important when one investigates the response of neurons to external inputs or network effects. In the later case it is important to take into account that the synaptic effective decay times of excitation and inhibition are affected by the rescaling parameters.

## Acknowledgments

This work was supported by NSF grants DMS-1313861 (HGR) and DMS-1608077 (HGR).

